# Selective targeting of an oncogenic *KRAS* mutant allele by CRISPR/Cas9 induces efficient tumor regression

**DOI:** 10.1101/807578

**Authors:** Qianqian Gao, Wenjie Ouyang, Bin Kang, Xu Han, Ying Xiong, Renpeng Ding, Yijian Li, Fei Wang, Lei Huang, Lei Chen, Dan Wang, Xuan Dong, Zhao Zhang, Yanshan Li, Baichen Ze, Yong Hou, Huanming Yang, Yuanyuan Ma, Ying Gu, Cheng-chi Chao

**Affiliations:** Guangdong Provincial Key Laboratory of Genome Read and Write, BGI-Shenzhen, Shenzhen 518083, China; China National GeneBank, BGI-Shenzhen, Jinsha Road, Shenzhen 518120, China; Guangdong Provincial Academician Workstation of BGI Synthetic Genomics, BGI-Shenzhen, Shenzhen, 518083, China; Department of Thoracic Surgery II, Key Laboratory of Carcinogenesis and Translational Research (Ministry of Education/Beijing), Peking University Cancer Hospital and Institute, Beijing, China; BGI Education Center, University of Chinese Academy of Sciences, Shenzhen, 518083, China; James D. Watson Institute of Genome Sciences, Hangzhou 310058, China; Ab Vision, Inc, Milpitas, California, USA

**Keywords:** *KRAS*, CRISPR-Cas9, dCas9-KRAB, gene-editing, mRNA-regulating, oncogenic mutation, bioinformatic pipeline

## Abstract

**Background:** *KRAS* is one of the most frequently mutated oncogenes in human cancers, but its activating mutations have remained undruggable due to its picomolar affinity for GTP/GDP and its smooth protein structure resulting in the absence of known allosteric regulatory sites.

**Results:** With the goal of treating mutated *KRAS*-driven cancers, two CRISPR systems, CRISPR-SpCas9 genome-editing system and transcription-regulating system dCas9-KRAB, were developed to directly deplete *KRAS* mutant allele or to repress its transcription in cancer cells, respectively, through guide RNA specifically targeting the mutant but not wild-type allele. The effect of *in vitro* proliferation and cell cycle on cancer cells as well as *in vivo* tumor growth was examined after delivery of Cas9 system. SpCas9 and dCas9-KRAB systems with sgRNA targeting the mutant allele both blocked the expression of mutant *KRAS* gene, leading to an inhibition of cancer cell proliferation. Local adenoviral injections using SpCas9 and dCas9-KRAB systems both suppressed tumor growth *in vivo*. The gene-depletion system (SpCas9) performed more effectively than the transcription-suppressing system (dCas9-KRAB) on tumor inhibition. Application of both Cas9 systems to wild-type *KRAS* tumor cells did not affect cell proliferation *in vitro* and *in vivo*. Furthermore, through bioinformatic analysis of 31555 SNP mutations of the top 20 cancer driver genes, we showed that our mutant-specific editing strategy could be extended to a list of oncogenic mutations with high editing potentials, and this pipeline can be applied to analyze the distribution of PAM sequence in the genome to survey the best targets for other editing purpose.

**Conclusions:** We successfully developed both gene-depletion and transcription-suppressing systems to specifically target an oncogenic mutant allele of *KRAS* which led to significant tumor regression. It provides a promising strategy for the treatment of tumors with driver gene mutations.

## Background

High frequency of *RAS* mutations has been found in various types of human cancers, including colon^1,2^, lung^3^ and pancreatic^4^ cancers which are the most deadly malignancies worldwide^5^. The three *RAS* oncogenes including *NRAS*, *HRAS* and *KRAS* make up the most frequently mutated gene family in human cancers. *KRAS* mutation is the most prevalent (21%) among the three genes, while the other two are 3% and 8% for *NRAS* and *HRAS*, respectively^6^.

*KRAS* is predominantly mutated in pancreatic ductal adenocarcinomas (PDACs), colorectal adenocarcinomas (CRCs) and lung adenocarcinomas (LACs)^7^. Majority of oncogenic *KRAS* mutations occur at codon 12, 13, and 61. G12 mutations are the most common variations (83%). It was reported that *KRAS* G12S is present in 1.84% of all colorectal adenocarcinoma patients, while in non-small cell lung carcinoma the frequency is 0.5%^8^ (Table 1).

**Table 1.**
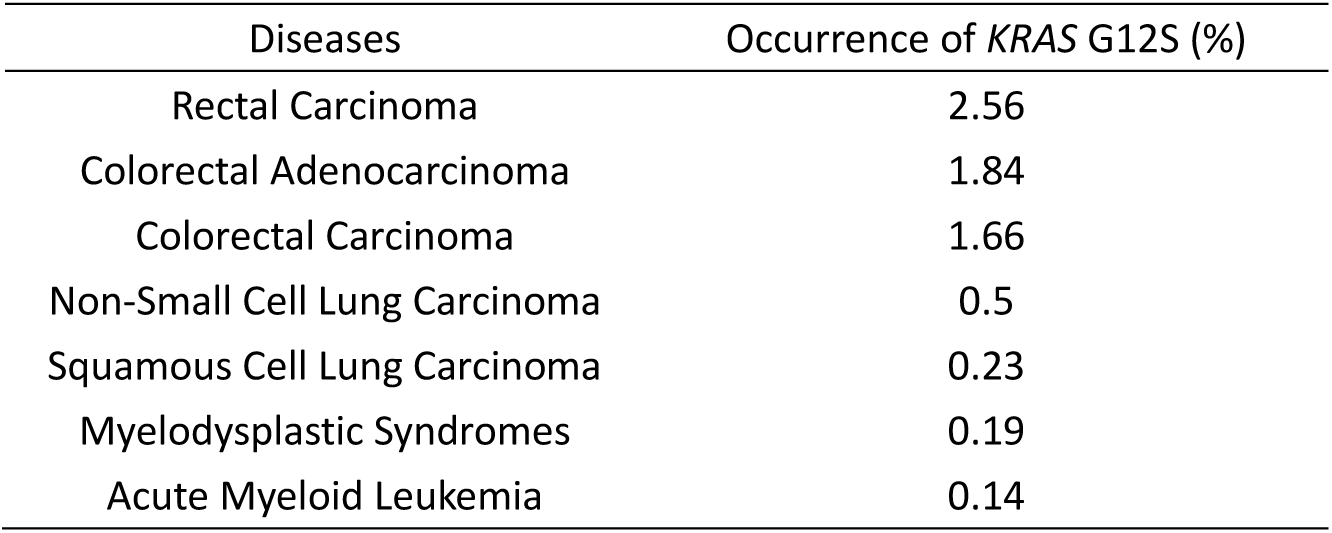
Occurrence of *KRAS* G12S mutation in different diseases

| Diseases | Occurrence of *KRAS* G12S (%) |
| --- | --- |
| Rectal Carcinoma | 2.56 |
| Colorectal Adenocarcinoma | 1.84 |
| Colorectal Carcinoma | 1.66 |
| Non-Small Cell Lung Carcinoma | 0.5 |
| Squamous Cell Lung Carcinoma | 0.23 |
| Myelodysplastic Syndromes | 0.19 |
| Acute Myeloid Leukemia | 0.14 |

Comprehensive efforts have been stimulated to develop therapeutic strategies to halt mutant *KRAS* function for cancer treatment, based on the well validated role of mutation-induced activation of KRAS in driving cancer development and growth. Different strategies to inhibit *KRAS* signaling have been under investigation, including exploring direct KRAS-binding molecules, targeting proteins that facilitate KRAS membrane-associated or downstream signaling, searching for synthetic lethal interactors, and novel ways of inhibiting *KRAS* gene expression and harnessing the immune system^9,10^. However, after more than three decades of research efforts, anti-KRAS therapy has not shown an effective clinical benefit.

The numerous studies to block *RAS* pathway have demonstrated the necessity to pursue mutation-specific *RAS*-targeted strategies. Small molecules that selectively bind to the *KRAS* G12C mutant were reported but limitedly demonstrated *in vitro*^11^. Gray *et al*. have also targeted *KRAS*-G12C by a GDP analogue which could covalently bind to the cysteine of G12C mutant, yet with a limitation to penetrate into cells^12^. Synthetic lethal interactors have also been screened in G13D^13,14^ or Q61K^15^ mutant cell lines to specifically target cancer cells, but with a far distance to be clinically applied. Despite the various attempts to directly interfere KRAS published previously, KRAS protein with a structure lacking suitable binding pocket for small molecule inhibitors, still remains a challenging target for therapeutic purpose^10^.

Development of antibodies and small molecule inhibitors is cost-ineffective and time consuming. Compared to the traditional antibody or inhibitor which can only be used to alter one specific target, genome editing technology could be a better alternative to flexibly manipulate biological activity of designated molecules at DNA level. CRISPR (Clustered regularly interspaced short palindromic repeats)/SpCas9 (CRISPR associated protein 9) system, developed from *Streptococcus pyogenes*, recognizes specific DNA sequences and is widely applied to genome editing of mammalian cells^16,17^. Taeyoung Koo *et al*. has used CRISPR/Cas9 to target an epidermal growth factor receptor (*EGFR*) oncogene harboring a single-nucleotide missense mutation to enhance cancer cell killing^18^, while Zhang-Hui Chen *et al*. targeted genomic rearrangements in tumor cells through insertion of a suicide gene by Cas9^19^. Those findings have proved the concept of specifically disrupting mutant tumor cells by CRISPR/Cas9. *KRAS* mutant alleles including G12V, G12D and G13D, have also been targeted by CRISPR/Cas9 system to control tumor growth^20,21^. In addition, CRISPR-Cas13a system was engineered for targeted therapy of *KRAS*-G12D and *KRAS*-G12C mutants in pancreatic cancer^22^. The above three *KRAS* mutant alleles become druggable by using CRISPR/Cas9 genome-editing system. However, G12S mutation, a prevalent mutation in colorectal adenocarcinoma, has not been targeted by CRISPR system until now.

Here we demonstrated G12S mutant allele, but not the wild type *KRAS* can be specifically targeted by CRISPR-SpCas9 system. The delivery of SpCas9 and a guide RNA targeting G12S mutant allele in cancer cells affected the *in vitro* proliferative ability and cell cycle of tumor cells, and the *in vivo* tumor growth. Besides genome-editing CRISPR-SpCas9 system, transcription-regulating dCas9-KRAB (dead Cas9, dCas9; the Krüppel associated box, KRAB) system, which binds to target sequence by dCas9 and downregulate mRNA transcription by transcriptional repressor KRAB, was applied to inhibit tumor growth, but the effectiveness of dCas9-KRAB was not comparable to the genome-editing system. Additionally, the specific CRISPR targeting sites of 31555 oncogenic mutations in top 20 cancer driver genes were screened using our high-throughput bioinformatics analysis, which facilitated the application of genome editing strategy to other cancer mutations. To the best of our knowledge, our study is the first to target *KRAS*-G12S mutant by CRISPR/Cas9 and dCas9-KRAB systems for inhibition of tumor growth. Moreover, our bioinformatic pipeline for analyzing the distribution of protospacer adjacent motif (PAM) sequence provided a useful tool for editing targets screening. Combined with next generation sequencing (NGS), the genome-editing approach would be a promising strategy for targeting *KRAS* or other oncogenic mutations for personalized cancer treatment.

## Methods

### Cell lines and cell culture

HEK293T cells (ATCC, CRL-11268) were purchased from the American Type Culture Collection (ATCC). HEK293T cells were cultured in Dulbecco’s modified Eagle’s medium (DMEM, Gibco, 21063029) supplemented with 10% fetal bovine serum (Hyclone, SH30084.03HI), penicillin (100 IU/ml), and streptomycin (50 µg/ml). A549 and H2228 cell lines were purchased from Shanghai Cellbank, China. And they were cultured in RPMI-1640 medium (Gibco, C22400500BT) supplemented with 10% fetal bovine serum (Hyclone, SH30084.03HI), penicillin (100 IU/ml), and streptomycin (50 µg/ml).

### Plasmid construction

pX330-U6-Chimeric vector (Addgene, #42230) and lentiCRISPR v2 plasmidwith puromycin-resistance (Addgene, #52961) were purchased from Addgene. For sgRNA expression, oligonucleotides containing each target sequence were synthesized (BGI), followed by annealing in a thermocycler. Annealed oligonucleotides were ligated into the lentiCRISPR v2 plasmid digested with Bsm BI (Supplemental Figure S1).

### Lentivirus production

HEK293T cells were seeded at 70-80% confluency on 100 mm dishes. One day after seeding, the cells were transfected with a mixture (18 µg) of transfer plasmid (empty lentiCRISPR v2 or lentiCRISPR v2 containing sgRNA), psPAX2 (Addgene, #12260), and pMD2.G (Addgene, #12259) at a weight ratio of 4:3:2 using 54 µL PEI (Polysciences, 24765-1, 1 µg/µl). We changed the medium after 4-6 hours of incubation at 37 °C and 5% CO_2_. Viral supernatants were collected 72 hours after transfection and filtered through a 0.45 μm filter (Millipore, SLHP033RB), and ultra-centrifuged for 1.5 hours at 35,000 rpm (TYPE 45 Ti rotor of Beckman) at 4 °C to concentrate the virus. The resulting pellet was then resuspended in RPMI1640 medium without FBS, and stored at −80 °C. The lentiviral titers were determined with a Lenti-X™ qRT-PCR Titration Kit (Clontech).

### Cell transfection

A549 and H2228 cells were seeded at 70% confluency on six-well plate. One day after seeding, the cells were transfected with 3 µg target plasmid (empty lentiCRISPR v2 or lentiCRISPR v2 containing sgRNA), using 9 µL PEI. This medium was replaced with fresh culture medium 24 hours after transfection, and the cultures were supplemented with 2 μg/ml puromycin (InvivoGen, ant-pr) and incubated at 37 °C and 5% CO_2_.

### *In vitro* lentiviral transduction

For viral infection, A549 and H2228 cells were seeded into six-well plates at 1 × 10^5^ cells/well in the presence of 10 μg/ml polybrene and incubated with virus-containing medium. This medium was replaced with fresh culture medium 24 hours after infection, and the cultures were supplemented with 2 μg/ml puromycin (InvivoGen, ant-pr) and incubating for 48 hours. Subsequently, the double-transduced cells were counted and subjected to other assays.

### T7E1 assay

Genomic DNA was isolated using the Genomic DNA Kit (Tiangen, #DP304-03) according to the manufacturer’s instructions. The region of DNA containing the nuclease target site was amplified by PCR with the following primers: *KRAS* forward, 5’-atgcatttttcttaagcgtcgatgg-3’; *KRAS* reverse, 5’-ccctgacatactcccaaggaaag-3’. The PCR amplification was performed according to the following protocol: 2 min at 94 °C; 30 cycles of (10 s at 98 °C, 30 s at 56 °C, 25 s at 68 °C). After separation on a 2% agarose gel, size-selected products were purified using QIAquick Gel Extraction Kit (QIAGEN, 28706). The purified PCR products were denatured by heating and annealed to form heteroduplex DNA, and then treated with 5 units of T7 endonuclease 1 (New England Biolabs) for 30 min at 37°C and finally analyzed by 2% agarose gel electrophoresis.

### RNA extraction and qPCR

Total RNA was isolated from cells using TRIzol LS reagent (Invitrogen, 10296028) following the manufacturer’s protocol. One microgram of RNA was then reverse transcribed using Primescript RT Reagent (Takara, RR047A). Quantitative PCR was performed using Fast Sybr Green Master mix (Applied Biosystems) and the primers were: *KRAS* forward, 5’-atgcatttttcttaagcgtcgatgg-3’; *KRAS* reverse, 5’-ccctgacatactcccaaggaaag-3’. Each messenger RNA (mRNA) level was measured as a fluorescent signal normalized based on the signal for glyceraldehyde 3-phosphate dehydrogenase (GAPDH). Relative quantification was determined by the ^ΔΔ^Ct method and normalized according to GAPDH.

### Cell proliferation assay and cell cycle analysis

Cells were seeded in 96-well plates at 1 × 10^3^ per well in 90 µL cell medium. Cell proliferation was accessed by Cell Counting Kit-8 (YEASEN, 40203ES80) according to the manufacturer’s instructions. Briefly, 10 µL of CCK-8 solution was added to cell culture and incubated for 3-4 hours. Cell proliferation was evaluated according to the absorbance at 450 nm wave length. For analyzing cell cycle, cells were plated in six-well plates at 6 × 10^5^ per well. After staining by propidium iodide (Sigma–Aldrich), the cell cycle distribution was analyzed by flow cytometry.

### Colony formation assay

A549 and H2228 cells were plated in six-well plates at 2 x 10^2^ per well and maintained in RPMI1640 medium supplemented 10% FBS. After 2 weeks, the cells were washed once with PBS, fixed with cold methanol for 10 min, and then stained with 0.5% Crystal violet. The number of colonies was calculated by ImageJ. All these experiments were performed in triplicates.

### Western blot analysis

A549 and H2228 cells were plated in six-well plates at a confluent of 70%. 48 hours after adenovirus infection, whole-cell extracts were prepared by lysing cells with adding 500 µL hot SDS-PAGE buffer (Beyotime, P0015B). Tumor tissues were homogenized by TGrinder (Tiangen, OSE-Y30), and lysed with RIPA buffer containing complete protease inhibitor cocktail (Roche). Target proteins were detected by western blot analysis with the following antibodies: GAPDH mouse monoclonal antibody (Proteintech, 60004-1-Ig), Akt (pan) (40D4) mouse monoclonal antibody (Cell Signaling, 2920), Phospho-Akt (Ser473) (D9E) XP Rabbit mAb (Cell Signaling, 4060), p44/42 MAPK (Erk1/2) (137F5) Rabbit mAb (Cell Signaling, 4695), Phospho-p44/42 MAPK (Erk1/2) (Thr202/Tyr204) (D13.14.4E) (Cell Signaling, 4370), mouse monoclonal Anti-MAP Kinase, activated (Diphosphorylated ERK-1&2) antibody (Sigma, M8159), Ras Antibody (Cell Signaling, #3965) and Anti-RAS (G12S) Mouse Monoclonal Antibody (NewEast, 26186).

### Generation, treatment and analysis of tumor xenografted mice

Xenograft mouse model of human lung cancer tumors were implanted under the left upper limb in the abdomens of 6- to 8-week old male NCG mice by subcutaneous injection of A549 (5×10^6^ cells in 200 µL DPBS (Gibco, C14190500BT)) or H2228 cells (2× 10^6^ cells in 200 µL DPBS). After tumor cell injection, when tumor volumes reached a range of 50–100 mm^3^, mice were randomly separated to one of five groups to receive PBS, AdV-Cas9, AdV-Cas9-sgG12S, Lenti-v2, or dCas9-KRAB-sgG12S (nine mice per group). The first day of treatment was designated as day 1. PBS, Adenovirus (1 × 10^9^ PFU in 10 µL DPBS), or lentivirus (5 × 10^10^ copies in 70 µL DPBS) was administered intratumorally on day 1, 4 and 7. Tumor growth inhibition was evaluated twice a week by measuring the length (L) and width (w) of the tumor. Tumor volume was determined using the following formula: volume = 0.523L(w)^2^.

### H&E staining

Formalin-fixed and paraffin-embedded tumor tissues were cut into sections and stained with hematoxylin and eosin (H&E). Histopathology was reviewed by an experienced pathologist.

### IHC staining

Tumor tissues were formalin-fixed, paraffin-embedded and stained using anti-RAS (G12S) mouse monoclonal antibody (NewEast, 26186) followed by incubation with HRP-conjugated corresponding secondary antibody (Sigma-Aldrich). The expression levels were evaluated by H-score method. Scoring was independently reviewed in parallel by two experienced pathologists.

### Analysis of off-target effects

Paired-end reads of each sample were aligned against the sequence of each off-target locus (~150bp) using BWA-MEM^23^ (version 0.7). The mapped reads for each off-target locus were then obtained from the alignment result. Mapped reads number (M) for each off-target locus could be got by using SAMtools idxstats module^24,25^. By applying a tool called FLASH^26^, the mapped paired-end reads were merged. By using regular expression to search the sequences of off-targets’ protospacers and their reverse complementarity sequence in the above merged read files, the number of protospacers (S) among the mapped reads could be obtained. The editing efficiency for an off-target could be obtained via the following equation:

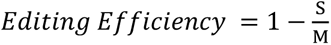

### PAM analysis

Annotate and prioritize genomic variants based on previous report^27^. Briefly, use ANNOVAR^28^ to annotate COSMIC v88 mutation database (perl table_annovar.pl humandb/hg19_cosmic88.txt humandb/ -buildver hg19 -out cosmic -remove -protocol refGene -operation gx -nastring . -csvout), and select variants located in the exons of the 20 cancer driver genes. Based on the gene mutation and wild-type genome information (https://www.ncbi.nlm.nih.gov/assembly/GCF_000001405.25), we applied Pandas (https://pandas.pydata.org/), a python package, to analyze the COSMIC SNP mutation information to generate a data frame. We applied Pyfaidx^29^, a python package to extract specific sequences from the GRCh37.p13 reference genome. PAM sequences of SpCas9, SaCas9, and LbCpf1 CRISPR nucleases were analyzed in the GRCh37.p13 reference genome. Once the SNP mutation is in the seed region of PAM sequences, we consider it can be edited by CRISPR nucleases.

### Statistical analysis

Significance of all data was determined using two-tailed Student’s t-test, and p-values <0.05 were considered statistically significant.

## Results

### Cas9-sgG12S specifically targets *KRAS* mutant alleles

*KRAS* gene locates in the short arm of human chromosome 12. There are four dominant mutant alleles at G12 position in exon 1, G12S (c.34G>A), G12V (c.35G>T), G12C (c.34G>T) and G12D (c.35G>A) (Figure 1A). These single nucleotide missense mutations are next to a PAM (TGG) sequence recognized by SpCas9. Since variations of DNA base in the PAM or seed sequence affect the recognition of SpCas9, five sgRNAs in total were designed to target the four *KRAS* mutant alleles, including G12S (sgG12S), G12V (sgG12V), G12C (sgG12C) and G12D (sgG12D), and the *KRAS*-WT gene (single guide G12-wild type RNA, sgG12-WT).

**Figure 1.**
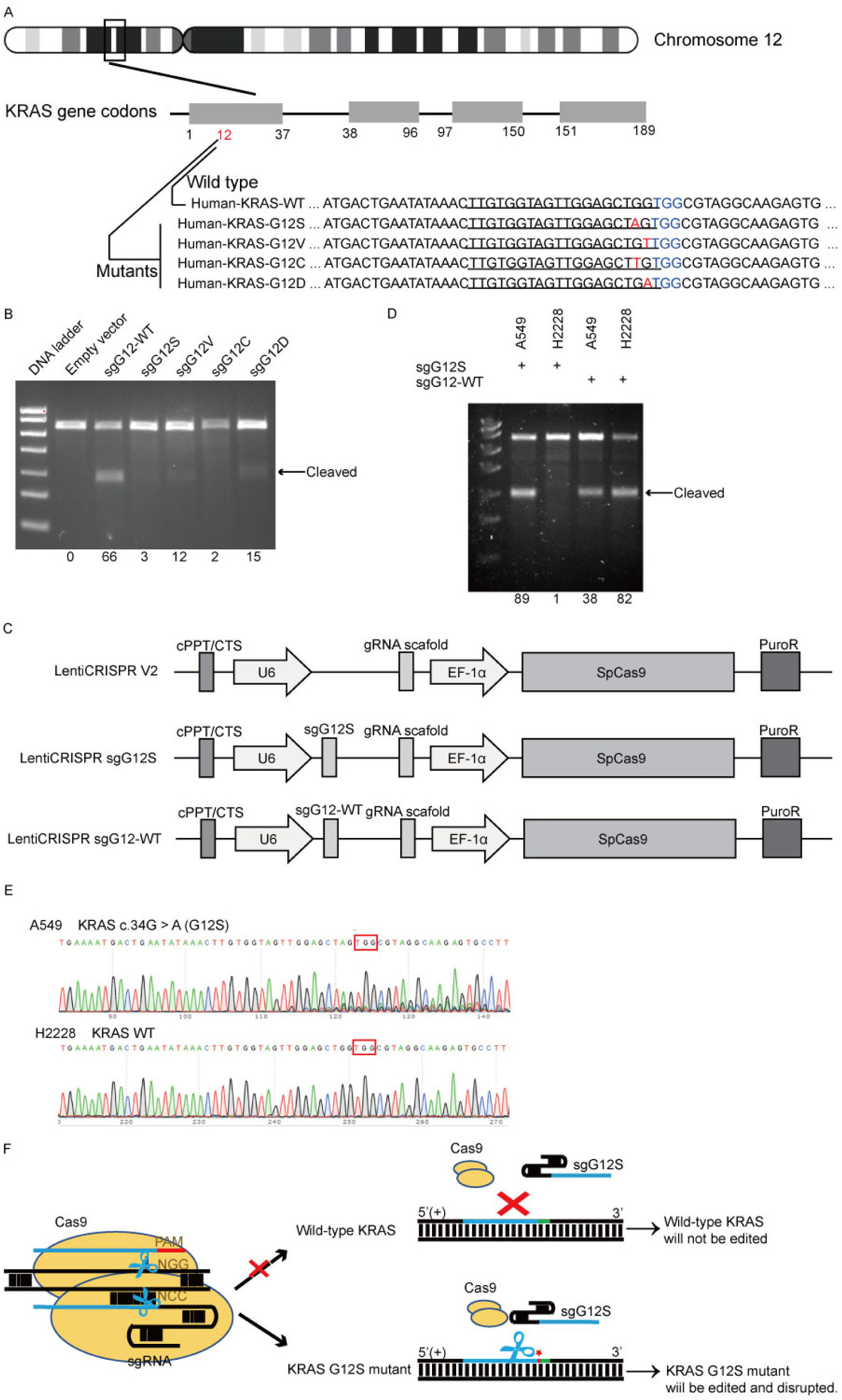
*KRAS* G12S oncogenic mutant-specific Cas9. **a** Mutations (red) at *KRAS* G12 site locate in the seed sequence of a PAM (blue). The human *KRAS* gene is located on chromosome 12. Oncogenic *KRAS* single-nucleotide substitutions within exon-1 of *KRAS* (c. 34G>A, c.35 G > T, c.34 G>T and c.35 G > A) result in G12S, G12V, G12C and G12D mutations. Design of their corresponding gRNAs was listed with a bottom line. **b** Editing efficiency of different gRNAs in 293T cells. Effective editing of genes is presenting by the appearance of cleaved band. And the gene editing efficiency is listed in lanes correspondingly. **c** Maps of lentiviral vectors, including LentiCRISPR V2 blank vector, sgG12S and WT guide RNA expressing vectors. **d** Efficiency and specificity of sgG12S and sgG12-WT in A549 and H2228 tumor cells infected with sgG12S or sgG12-WT lentiviruses 48 h post-infection. Effective editing of genes is presenting by the appearance of leaved band. And the gene editing efficiency is listed in lanes correspondingly. **e** Gene editing event was confirmed by sanger sequencing in A549 and H2228 cells. PAM sequence is marked in red box. **f** Diagram of the genome therapy strategy to target *KRAS* G12S mutant allele specifically. Blue strands: spacer; green strands: PAM sequence; red strands and star: single-nucleotide missense mutations.

We first examined the activity of these five sgRNAs in 293T cells (Figure 1B), which harboring the wild-type *KRAS* gene. To confirm the editing efficiency of sgG12-WT, and the specificity of sgG12-Mu (mutant), we transfected plasmids encoding spCas9 and different sgRNAs (Additional file 1: Figure S.1A) into 293T cells separately. We found that sgG12-WT disrupted *KRAS*-WT efficiently with an efficiency of 66% by T7E1 assay, while the editing efficiency of sgG12S, sgG12V, sgG12C and sgG12D in *KRAS*-WT were 3%, 12%, 2% and 15%, respectively (Figure 1B). Thus, sgG12S and sgG12C were more specific with much lower off-target effects on wildtype *KRAS*. Next, we confirmed the editing efficiency of sgG12S in A549 lung adenocarcinoma cells harboring *KRAS* G12S mutant allele. H2228, another lung adenocarcinoma cell line carrying no G12S mutant allele, was utilized as a negative control. Lentivirus containing spCas9-sgG12S or spCas9-sgG12-WT, and non-targeting control virus (Figure 1C) were respectively infected into A549 and H2228 cells. We found that spCas9-sgG12S edited *KRAS* G12S mutant allele in A549 cells with a high efficiency of 89%, while the editing efficiency was only 1% in wild-type *KRAS* allele in H2228 cells (Figure 1D). On the other hand, sgG12-WT edited *KRAS* in A549 and H2228 cells with editing efficiency of 38% and 82%, respectively, indicating that the sgG12-WT non-specifically bound to *KRAS* G12S sites with a high mismatch tolerance. To further confirm that sgG12S specifically edited *KRAS* G12S mutant allele, but not the wild-type allele, *KRAS* gene in puromycin selected A549 and H2228 cells was sequenced 2-3 days post infection (Figure 1E). *KRAS* in A549 was destroyed around PAM (TGG) sequence, while H2228 was not affected, further confirming the success of our spCas9-sgG12S system in efficient and specific targeting *KRAS* G12S allele (Figure 1F).

### Genome editing of *KRAS* G12S mutant allele inhibits the proliferation and cell cycle of tumor cell lines *in vitro*

To investigate whether targeting and disruption of the *KRAS* mutant allele by sgG12S could inhibit the proliferation of tumor cells, the cell numbers of A549 and H2228 cells were examined after gene editing (Figure 2A). The proliferation of sgG12S-targeted A549 cells was dramatically inhibited and almost retarded compared to non-targeting control and untreated groups. While the targeting of sgG12S had no effect on the proliferation of H2228 cells. Besides, a cell colony formation assay (CFA) (Figure 2B) and CCK-8 cell proliferation assay (Figure 2C) also confirmed the growth inhibition by Cas9-sgG12S targeting. As demonstrated in cell counting (Figure 2A), the proliferation of A549 cells was significantly suppressed shown in the CFA and CCK-8 assays. In contrast, the targeting of sgG12S had a less effect on the proliferation of H2228 cells carrying the wild-type *KRAS* allele.

**Figure 2.**
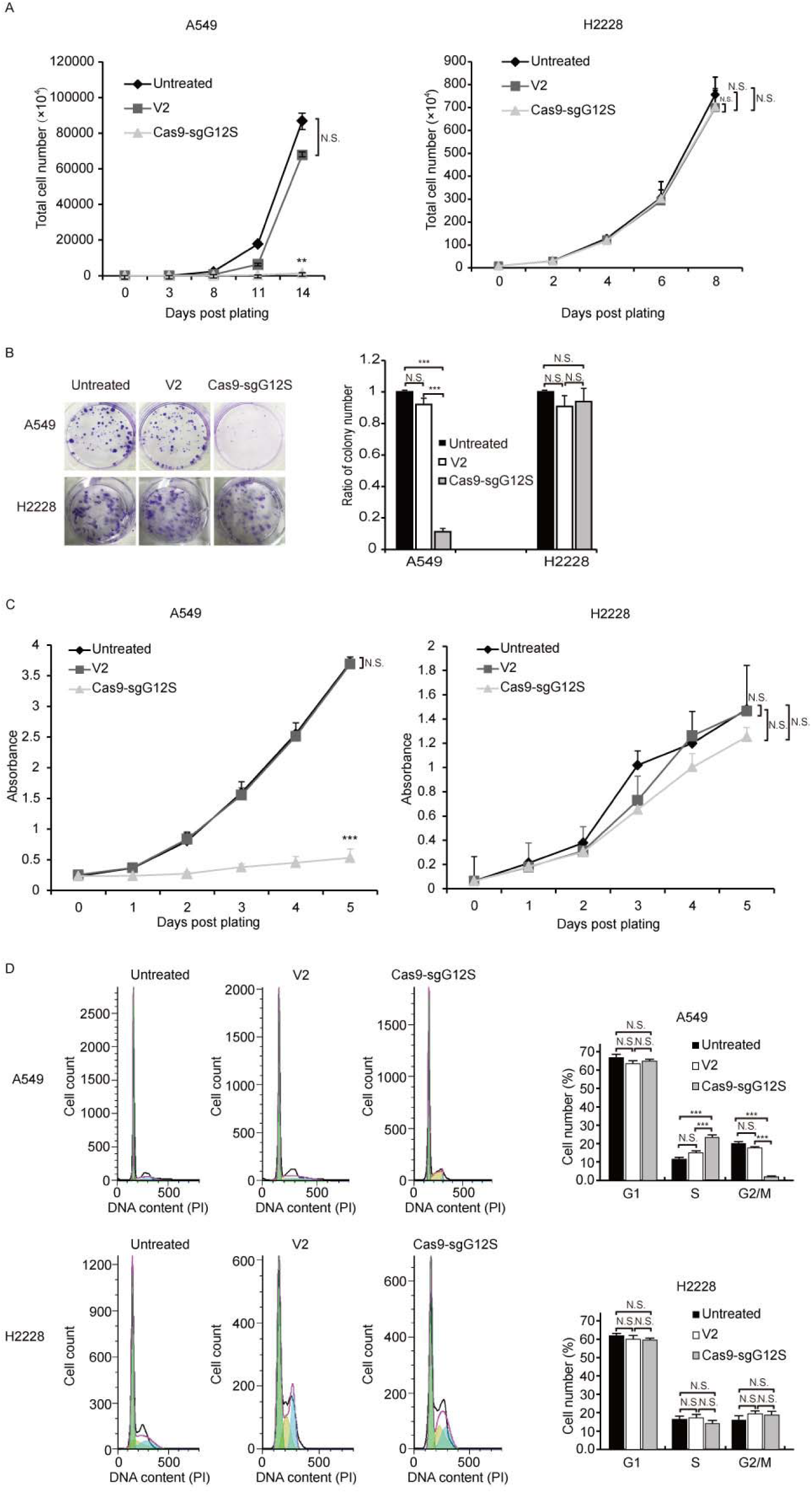
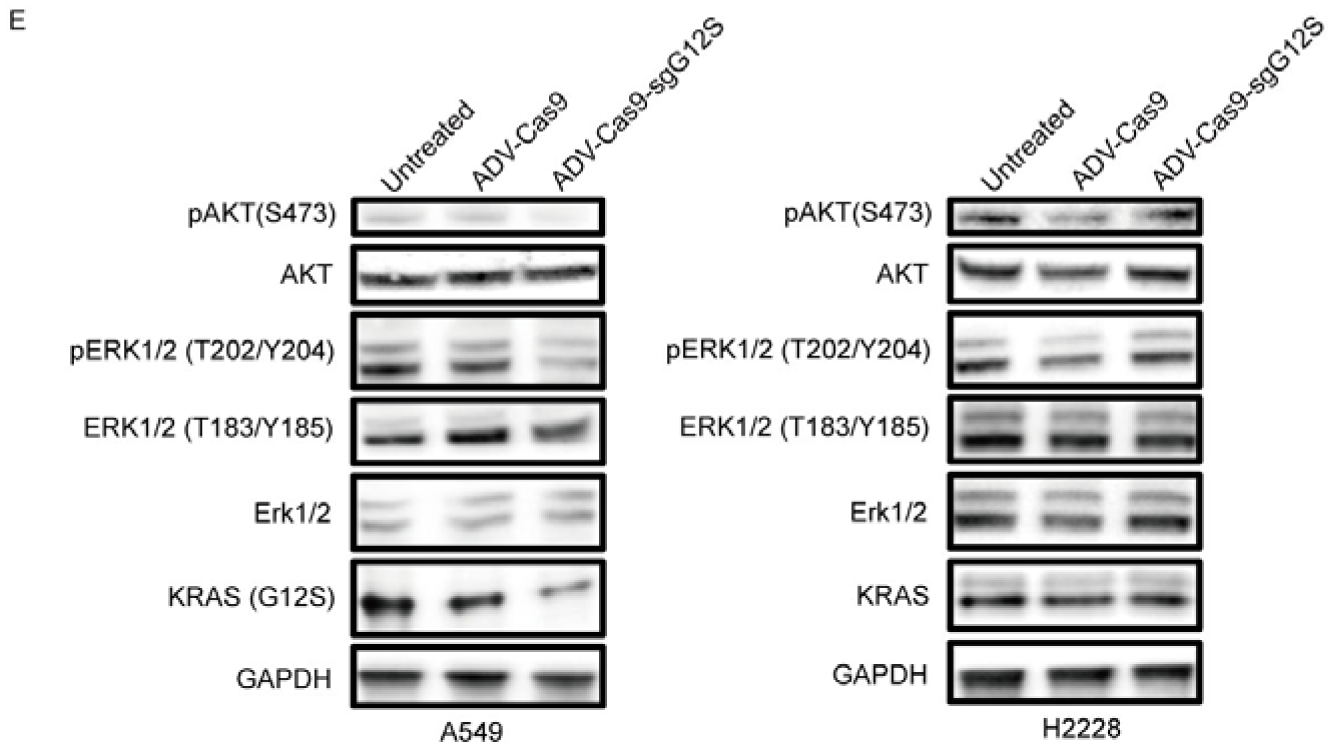
The anti-tumor effects of targeting *KRAS* G12S mutant allele *in vitro*. A549 and H2228 cells were subjected to cell proliferation (A), colony forming (B), CCK-8 (C), cell cycle (D) and WB (E) assays after treatment with lentiviral Cas9 and sgRNAs targeting *KRAS* G12S mutant allele. Error bars represent S.E.M. (∗) 0.01<P < 0.05, (∗∗) 0.001<P < 0.01, (∗∗∗) P < 0.001. **a** Cell growth curves determined by counting cell number with various treatments at different timepoints. **b** Colony formation assay in A549 and H2228 cells. Representative images of wells after 0.5% crystal violet staining are shown at left and colony number was determined 2 weeks after cell plating and treated with Cas9-sgG12S and sgG12-WT. **c** CCK-8 assay in A549 and H2228 cells. Cell proliferation was determined by use of CCK-8 reagents at different timepoints after plating. The number of cells in cultures with different treatments was accessed by the optical density at 490 nm of each CCK-8 reaction. **d** Cell cycle was determined by PI staining and FACS analysis. **e** Western blot analysis of the phosphorylation levels of AKT and ERK proteins.

We further assessed the cell cycle of sgG12S-targeted A549 and H2228 cells (Figure 2D). The Cas9-sgG12S treated A549 cells was mostly arrested at S phase, and the ratio of cell population at G2/M phase was downregulated correspondingly, while there was no effect on the cell cycle of sgG12S-treated H2228 cells. Next, we examined the activities of *KRAS* downstream signaling pathways including the expression and activation of AKT and ERK (Figure 2E). The treatment of Cas9-sgG12S in A549 tumor cells dramatically suppressed the expression of KRAS (G12S) protein, while the expression of wild-type KRAS protein in H2228 cells were not affected. Besides, the levels of phosphorylated-AKT (S473) and phosphorylated-ERK (T202/Y204) proteins were significantly downregulated in A549 cells edited with SpCas9-sgG12S, while another type of phosphorylated-ERK (T183/Y185) protein was not affected. As expected, AKT and ERK signaling pathways in H2228 cells were not affected by SpCas9-sgG12S. Collectively, our results suggested that the mutant allele-specific targeting by sgG12S can efficiently inhibit tumor cell proliferation and arrest the cycle of tumor cells at S phase, probably through downregulating AKT and ERK signaling pathways.

### Transcription-repressing system dCas9-KRAB inhibited proliferation of tumor cell lines *in vitro*

We next explored whether there were off-target effects of the mutant allele-specific nuclease outside of *KRAS* gene region by targeted deep sequencing at 14 potential off-target sites (Additional file 1: Supplemental Table 1). The potential off-target sites which were different from the on-target site by up to 4 nt mismatch in the human genome were identified by Feng Zhang lab’s CAS-OFFinder algorithm (http://www.rgenome.net/cas-offinder/). No indel was detected at these sites in Cas9-sgG12S treated A549 and H2228 tumor cells (Figure 3A, 3B).

**Figure 3.**
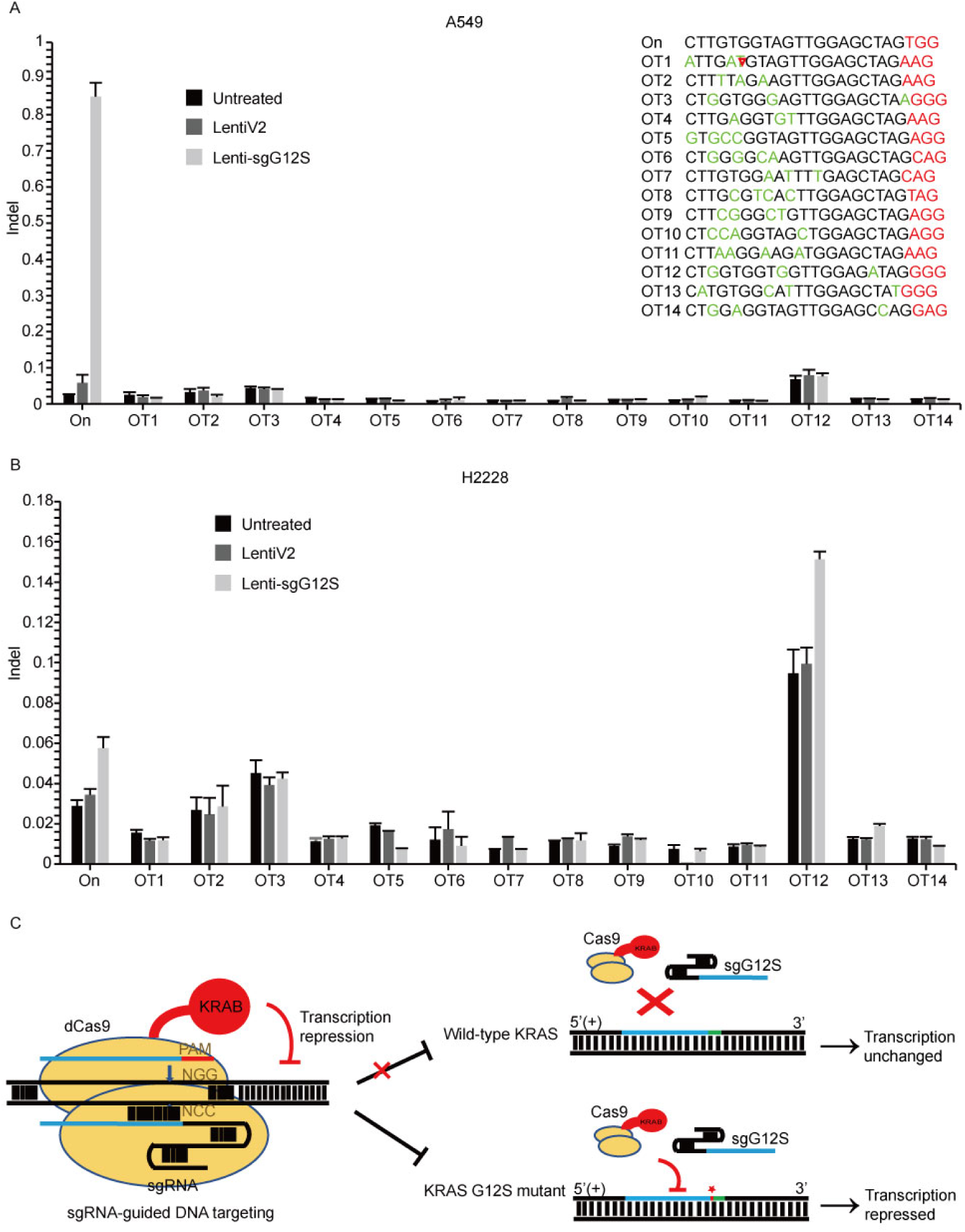
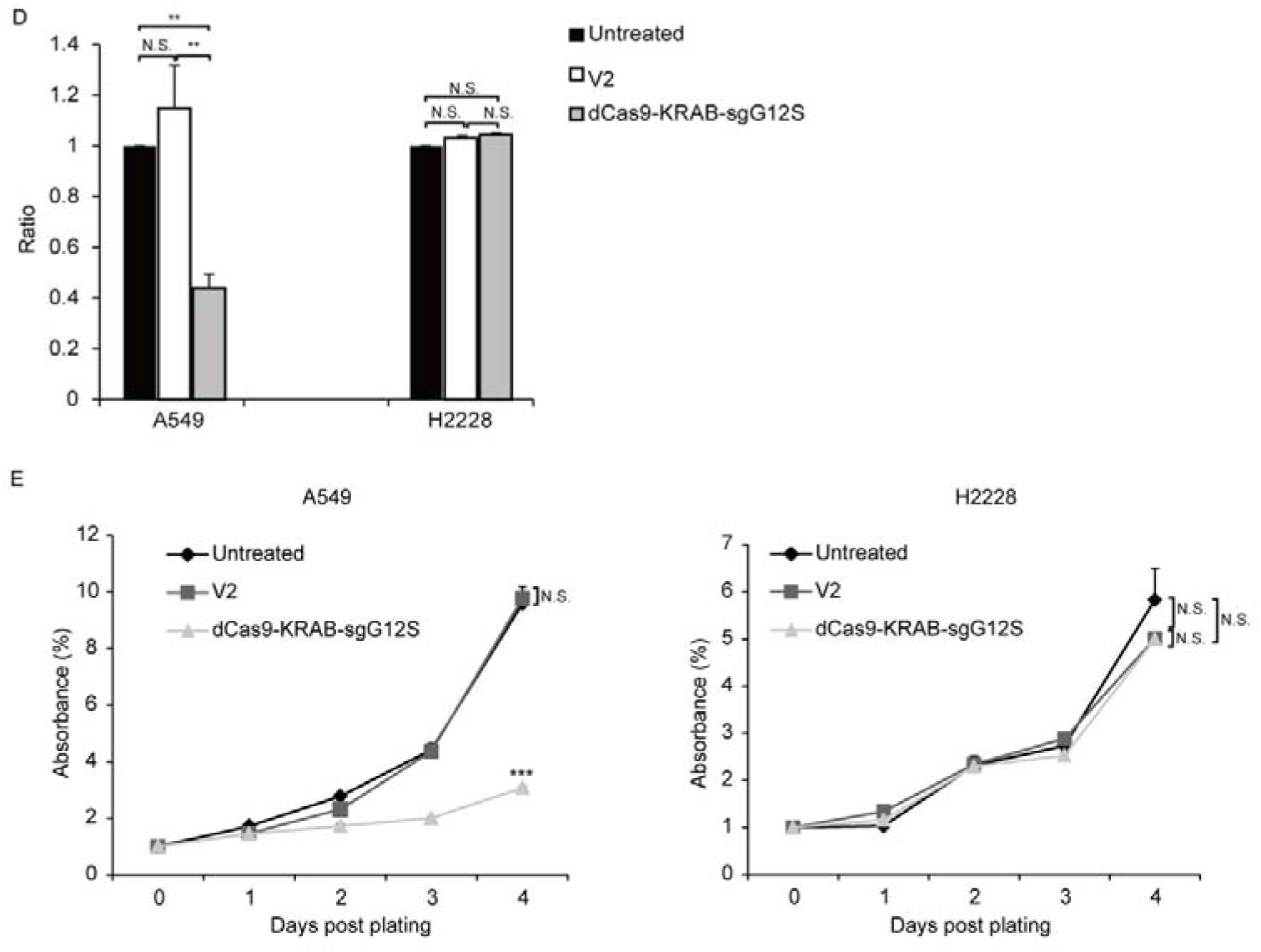
dCas9-KRAB mRNA-regulating system downregulated G12S transcription and inhibited tumor cell proliferation. **a, b** No off-target indels were detectably induced by CRISPR/Cas9 gene-cutting system at fourteen homologous sites that different from the on-target sites by up to 4 nt in the human genome. PAM sequences are shown in red and mismatched nucleotides are shown in green. On: on-target site. OT: off-target site. Cleavage position within the 20-bp target sequences is indicated by red arrow. Error bar indicates S.E.M. (n=3 to 4). **c** Diagram of knocking down *KRAS* G12S mutant allele specifically by dCas9-KRAB system. Blue strands: spacer; green strands: PAM sequence; red strands and star: single-nucleotide missense mutations. **d** qRT-PCR analysis of *KRAS* G12S mRNA expression. Error bars represent S.E.M. 0.01<P < 0.05, (∗∗) 0.001<P < 0.01, (∗∗∗) P < 0.001. **e** CCK-8 assay. Cell proliferation was determined by use of CCK-8 reagents at different timepoints after plating. The relative number of cells of each group with different treatments was determined by normalizing the optical density at 490 nm of each CCK-8 reaction to the average optical density of the negative control groups.

Genome-editing system has the likelihood to cause undesired double stand break (DSB) in the genome (Figure 1B, 1D). In order to avoid the undesired disruption of genome, we constructed a non-cutting transcription-regulating system, dCas9-KRAB system (Figure 3C), where KRAB is a transcriptional repressor to downregulate mRNA expression when binding to the regulatory elements of certain genes^30,31^. To test whether sgG12S linked to dCas9-KRAB may repress *KRAS* expression specifically in G12S mutant allele, A549 and H2228 cells were infected by dCas9-KRAB-sgG12S and non-targeting control lentivirus. As expected, the transcription of *KRAS* G12S mutant allele in dCas9-KRAB-sgG12S treated A549 cells was dramatically downregulated compared to non-targeting control or untreated cells (Figure 3D), while in H2228 cells, the transcription of wild-type *KRAS* was not affected in all three groups. In addition, the effect on tumor cell growth was also investigated by CCK-8 assay (Figure 3E). Consistently, the proliferation of dCas9-KRAB-sgG12S treated A549 cells was inhibited significantly compared to the controls, while no significant effect on H2228 tumor cell growth was observed. These results confirmed the *in vitro* specificity of the dCas9-KRAB system.

### Targeting *KRAS*-G12S mutant blocks tumor growth in tumor-bearing mice

To further explore the effects of *KRAS*-sgG12S targeting *in vivo*, AdV-Cas9-sgG12S and non-targeting control adenovirus were constructed and packaged (Additional file 1: Figure S2A). Lentivirus is relatively limited to use for *in vitro* or *ex vivo* gene delivery due to their restricted insertional capacities and relatively low titers.^32^ Therefore, the *in vivo* gene delivery experiments were conducted by adenoviral infection. The editing efficiency of AdVs was firstly confirmed in A549 and H2228 cells by T7E1 assay (Additional file 1: Figure S2B) and sanger sequencing (Additional file 1: Figure S2C). As expected, AdV-Cas9-sgG12S specifically edited *KRAS* G12S mutant allele in A549 cells, but not in H2228 cells harboring wild-type *KRAS* gene. In addition, AdV-Cas9-sgG12S inhibited the proliferation of A549, but not H2228 tumor cells *in vitro* (Additional file 1: Figure S2D).

Next, we examined the effect of sgG12S editing in cell-derived xenograft models of A549 and H2228 cells, respectively (Figure 4A-D). Local injection of AdV-Cas9-sgG12S significantly inhibited tumor growth, resulting in a 46% reduction in tumor volume (P<0.01) in A549-bearing mice (Figure 4A). In contrast, tumor volumes of control groups treated with either PBS or AdV-Cas9 vector grew over time, reaching an average size of more than 2000 mm^3^ 28 days after treatment (Figure 4A). As expected, no significant difference in tumor volume was showed in AdV-Cas9-sgG12S, AdV-Cas9 vector and PBS-treated mice implanted with H2228 cells containing the wild-type *KRAS* allele (Figure 4B). It confirmed the high specificity of *KRAS* G12S targeting *in vivo*. The tumor weight was also significantly decreased by 30% in animals treated with AdV-Cas9-sgG12S, compared to control groups treated with either AdV-Cas9 vector or PBS (P<0.05) in A549 bearing mice (Figure 4C). Consistent with tumor volume, there was no difference in tumor weight of H2228-implanted groups (Figure 4D).

**Figure 4.**
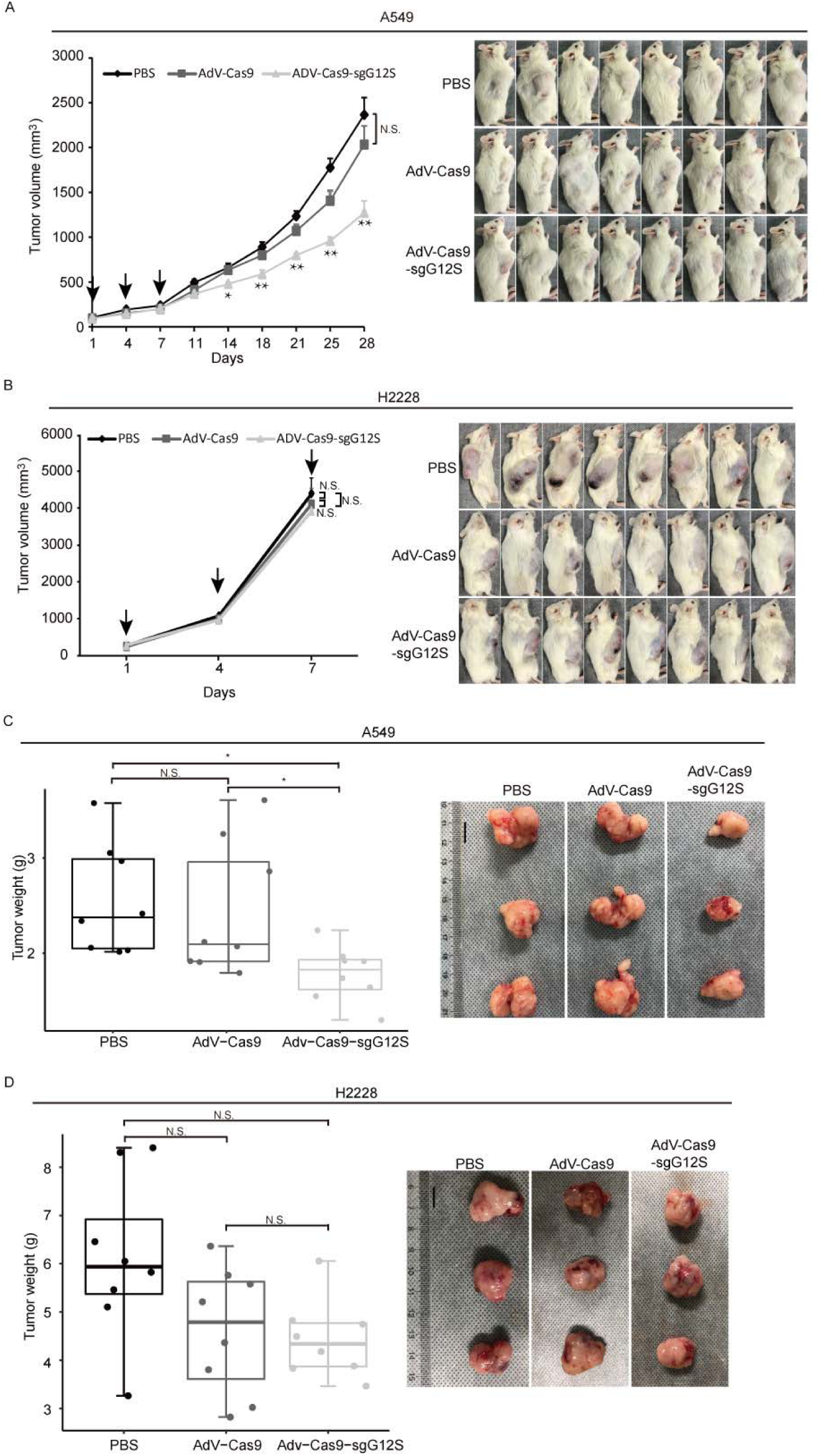
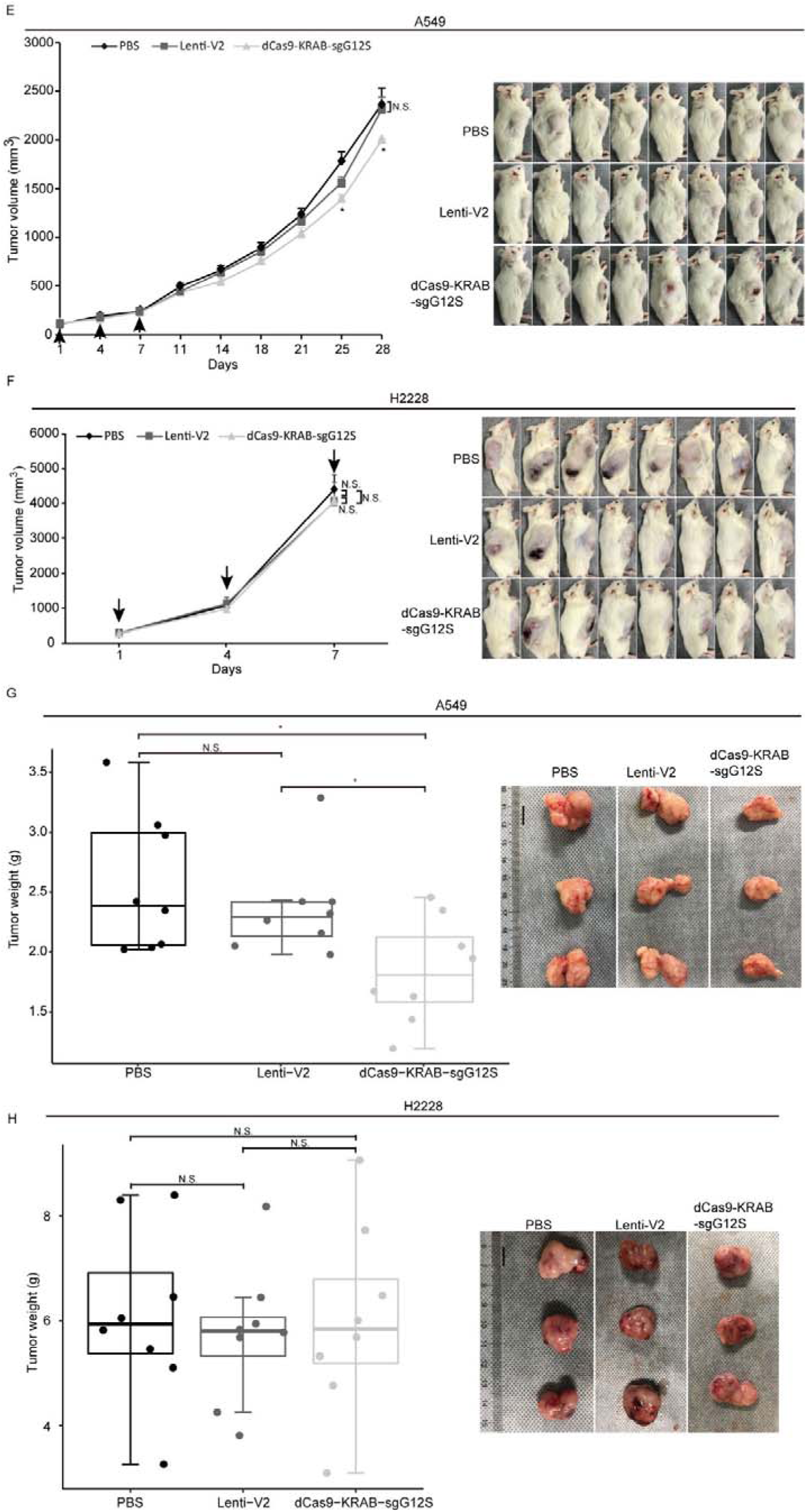
Antitumor effects of CRISPR-Cas9 and dCas9-KRAB systems in tumor xenograft models. Error bars represent SEM. 0.01<P < 0.05, (∗∗) 0.001<P < 0.01, (∗∗∗) P < 0.001. Values represent the mean ± S.E.M. (n=8 per group). **a, b** A549 and H2228 tumor-bearing mice were given intra-tumoral injections of PBS or AdV-Cas9 or AdV-Cas9-sgG12S adenoviruses on days 1, 4 and 7. Tumor growth was monitored twice a week post injection until tumor volume > 2000 mm^3^. **c**, **d** Weights of tumors removed from euthanized mice after 28 days in A549 tumor-bearing mice, and 7 days in H2228 tumor-bearing mice. **e**, **f** A549 and H2228 tumor-bearing mice were intra-tumoral injected of PBS or Lenti-V2 or dCas9-KRAB-sgG12S lentiviruses on day 1, 4 and 7. Tumor growth was monitored twice a week post injection until tumor volume > 2000 mm3. **g**, **h** Weights of tumors removed from euthanized mice after 28 days in A549 tumor-bearing mice, and 7 days in H2228 tumor-bearing mice.

To examine the efficacy of repressing G12S transcription by dCas9-KRAB system *in vivo*, NSG mice were xenografted with A549 and H2228 cells, and treated with dCas9-KRAB-sgG12S, non-targeting virus, or PBS when tumor size reached a volume of 100-200 mm^3^ (Figure 4E-H). The mice xenografted with A549 cells and treated with dCas9-KRAB-sgG12S showed 15.6% (P<0.05) decrease in tumor volume compared to a control (Figure 4E) and exhibited no notable metastasis or mortality during the observation period of 28 days. In contrast, the mice xenografted with H2228 cells treated with dCas9-KRAB-sgG12S did not show any inhibition of tumor growth but instead experienced a quick increase in tumor volume (Figure 4F). Similar rate of increase in tumor size also observed in mice treated with non-targeting vector or PBS. Tumor weights were also measured in mice treated with different viruses (Figure 4G, 4H). A significant decrease of tumor weight (28.2%, P<0.05) was observed in dCas9-KRAB-sgG12S treated mice xenografted with A549 cells (Figure 4G). In contrast, the H2228-bearing mice injected with either dCas9-KRAB-sgG12S, non-targeting vector or PBS treatment (Figure 4H) had little affected.

Throughout the mice study of gene-editing and transcription-repressing systems, no sign of weight loss (Additional file 1: Figure S3A-S3D) was observed. Taken together, these *in vivo* data suggested that gene targeting of mutant *KRAS* by SpCas9-sgG12S and dCas9-KRAB-sgG12S is effective and only restricted to the tumors with the *KRAS* mutations, with no obvious effects on the other cell types. Besides, CRISPR/Cas9 genome-editing system targeting mutant *KRAS* is more effective compared with the dCas9-KRAB mRNA-regulating system.

### Disruption of *KRAS*-G12S significantly inhibited the protein expression of the mutant KRAS in tumor-bearing mice

The antitumor efficacy of oncogenic mutant-specific gene-editing and mRNA-regulating systems were further investigated by western blot and immunohistochemical (IHC) staining in the xenograft tumor tissues disrupting *KRAS*-G12S mutant alleles (Figure 5). Western blot (WB) assay revealed a markedly reduced expression level of KRAS and KRAS G12S mutant proteins in the tumor tissues of A549 cells-engrafted mice edited by AdV-Cas9-sgG12S, but not in AdV-Cas9 treated control group. While in the tumor tissues of H2228 cells-engrafted mice, the expression level of wild-type KRAS protein was not dramatically changed in AdV-Cas9 or AdV-Cas9-sgG12S treated groups (Figure 5A). Consistently, dCas9-KRAB-sgG12S, but not V2 treated tumor tissues, exhibited markedly lower levels of both total and mutant KRAS proteins in A549-engrafted mice (Figure 5B). Importantly, tumor tissues from A549-engrafted mice treated with AdV-Cas9-sgG12S and dCas9-KRAB-sgG12S both showed significant reduction of KRAS G12S protein through in situ IHC staining, but such decrease was not observed in the control groups (Figure 5C, 5D). It implied that CRISPR/Cas9 system can efficiently target and reduce KRAS mutant protein expression. Taken together, these data indicate that the application of both the gene-cutting CRISPR-Cas9 and mRNA-regulating dCas9-KRAB systems could affect KRAS G12S protein downregulation *in vivo* and further result in a strong anti-tumor efficacy.

**Figure 5.**
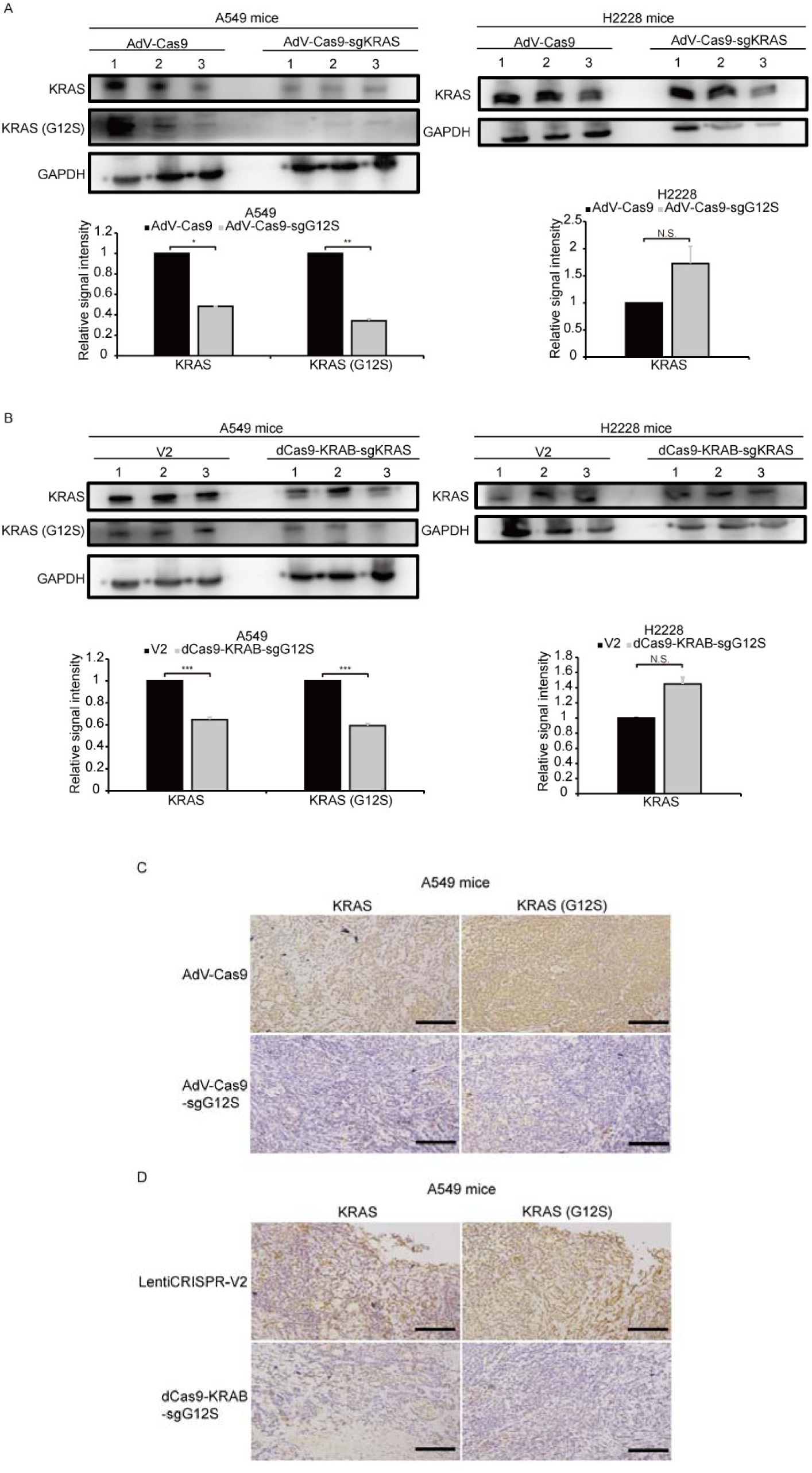
Targeting *KRAS* G12S mutant allele significantly inhibited the expression of KRAS mutant *in vivo*. Error bars represent SEM. (∗) 0.01< P < 0.05, (∗∗) 0.001< P < 0.01, (∗∗∗) P < 0.001. **a** Western blot analysis of the expression levels of total and mutant KRAS proteins in A549- and H2228-engrafted mice treated by CRISPR-Cas9 gene-editing system, respectively. The optical density analysis was performed from the results in three western blot replicate samples. Tumors were removed from euthanized mice after 28 days in A549 tumor-bearing mice, and 7 days in H2228 tumor-bearing mice. **b** Western blot analysis of the expression levels of total and mutant KRAS proteins in A549- and H2228-engrafted mice treated by dCas9-KRAB mRNA-regulating system, respectively. The optical density analysis was performed from the results in three western blot replicate samples. Tumors were removed from euthanized mice after 28 days in A549 tumor-bearing mice, and 7 days in H2228 tumor-bearing mice. **c** Immunohistochemical staining of KRAS and KRAS (G12S) were performed on tumor sections from A549 cells-engrafted mice treated with CRISPR-Cas9 gene-editing system. Scale bar: 100 µm. **d** Immunohistochemical staining of KRAS and KRAS (G12S) were performed on tumor sections from A549 cells-engrafted mice treated with dCas9-KRAB system. Scale bar: 100 µm.

### Extending the strategy of targeting tumor-specific mutant locus by gene editing system

Cas9-sgG12S editing system is a highly specific strategy to target cancer driver gene mutation with almost no difference in off-target effects in sgG12S and control groups in all cell lines we treated (Figure 3A, 3B). Moreover, Cas9-sgG12S targeting specifically and efficiently inhibits tumor growth, both *in vitro* and *in vivo*. Thus, this approach holds great potential to treat *KRAS* G12S mutation-driven cancers. In order to extend this strategy to different DNA nucleases to target other oncogenic mutations, driver gene mutations were collected from Cosmic database and the top 20 driver genes were selected to continue our proof-of-concept study (Figure 6A). These high-frequency driver gene mutations, including *JAK2*, *TP53*, *KRAS*, *EGFR*, etc., are widely spread in human malignancies (Figure 6B). Among these mutations, most of them are missense mutations, leading to single nucleotide variation (SNV) (Figure 6C). SNV occupies 74% of the whole mutations, while the percentage of deletion, insert and indel (insert and deletion) mutations is 16%, 7% and 3%, respectively.

**Figure 6.**
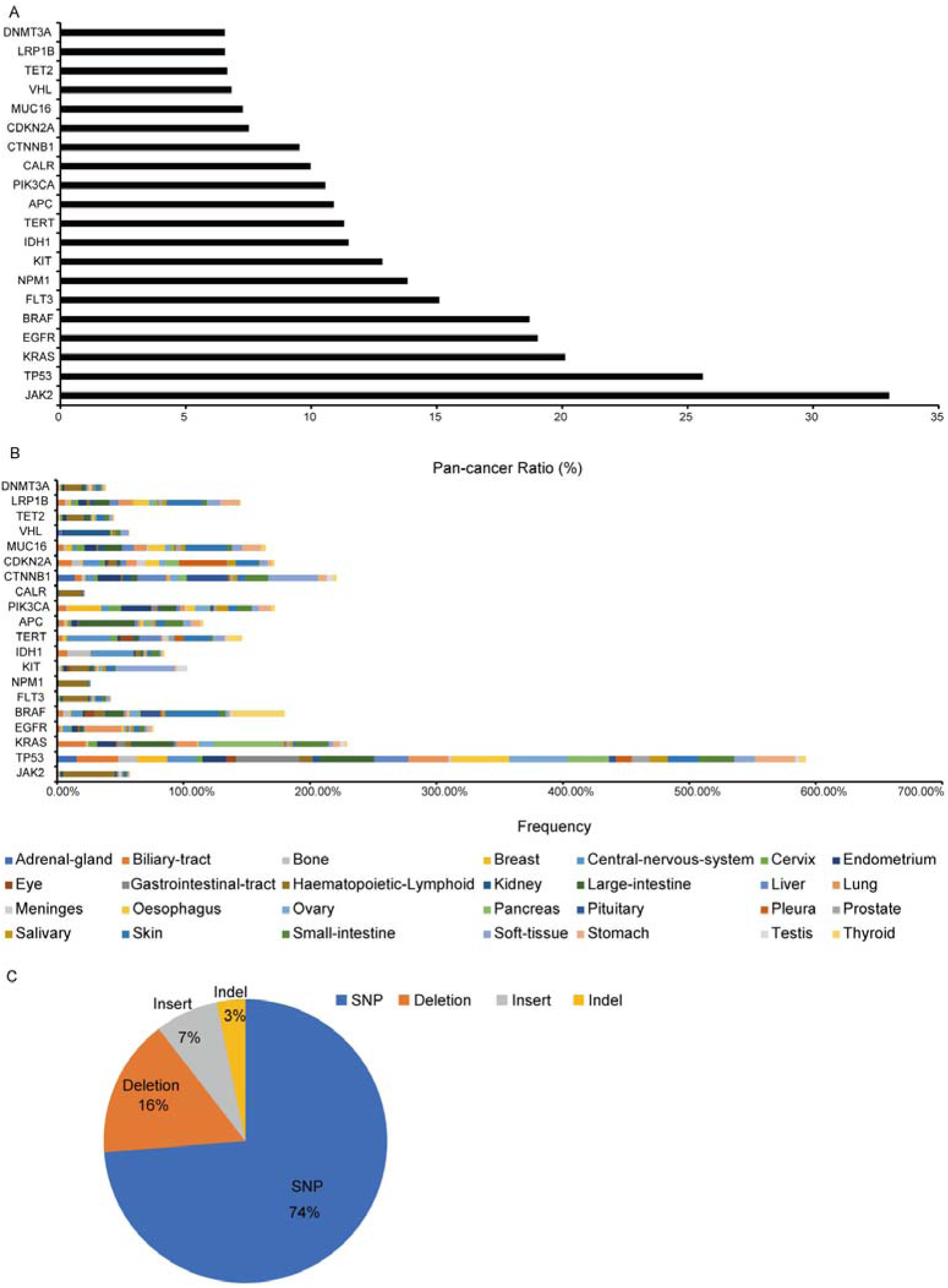
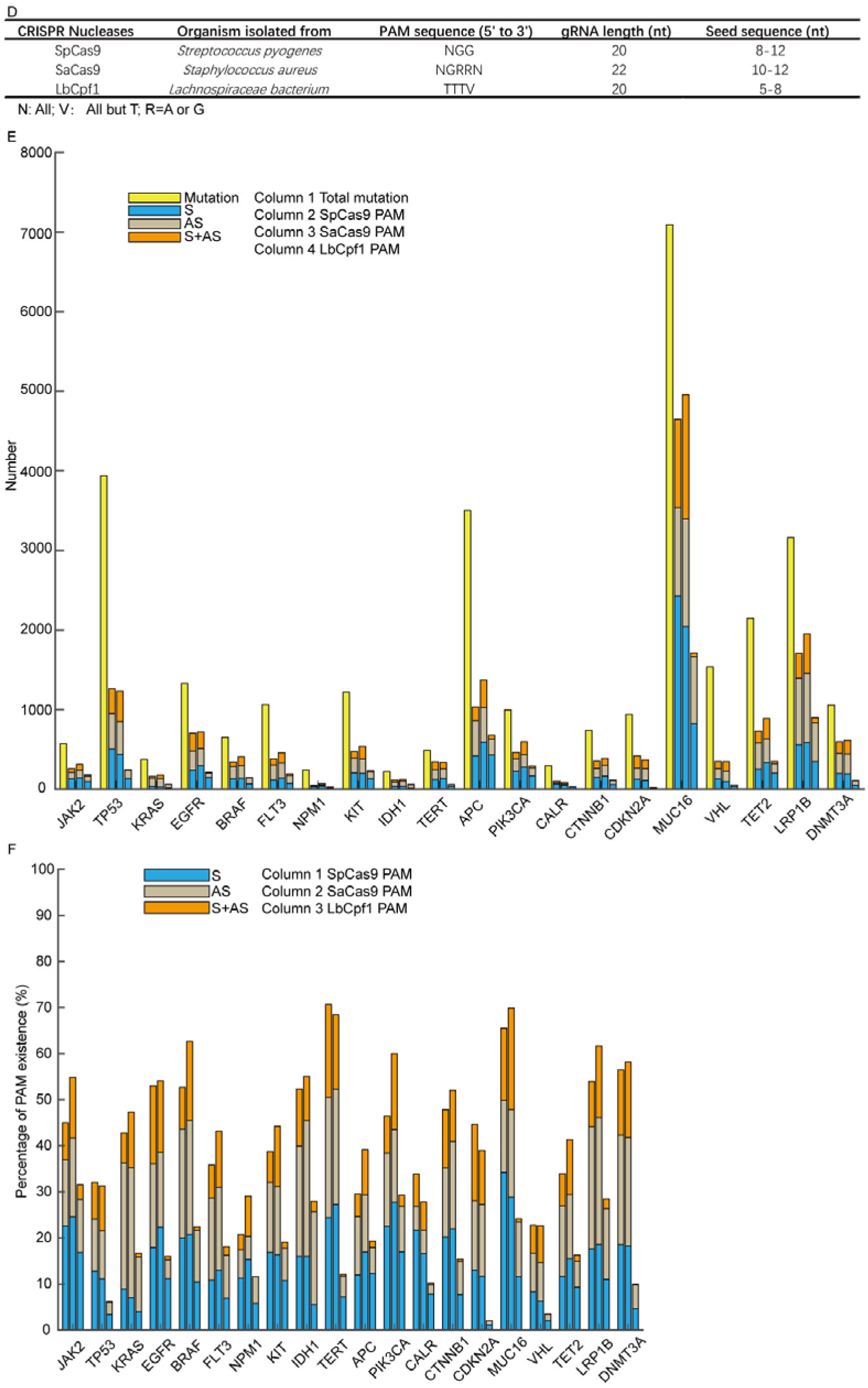
Screening of potential mutation-specific targets by CRISPR nucleases through bioinformatic analysis. **a** Top 20 oncogenic mutations discovered from Cosmic database. **b** Distribution of oncogenic mutations in human tissues. **c** Proportion of different mutation types of the top 20 oncogenic genes. **d** Characteristic of three mostly used CRISPR nucleases, SpCas9, SaCas9 and LbCpf1. **e** Statistics of mutations that are in seed sequences or PAM sequences. S, sense strand. AS, anti-sense strand. **f** Proportion of 31555 SNV oncogenic mutations that can be targeted by CRISPR nucleases. S, sense strand. AS, anti-sense strand.

There are large amounts of mutations of each cancer driver gene, and it is important to discover whether these oncogenic mutations can be edited or not and which DNA nucleases can be applied to edit them. To identify the mutations that could be specifically targeted by editing nucleases including SpCas9, SaCas9 and LbCpf1, we analyzed the SNV mutations to examine whether their flanking sequences fit the PAM or seed sequence requirements (Additional file 1: Figure S4). There is a length limitation of the seed sequence, and the seed sequence length of different nucleases is different (Figure 6D). In order to guarantee the targeting specificity, the lower limitation of the seed sequence length was used as threshold in our analysis (Additional file 1: Figure S4). Among the 31555 SNV mutations of the 20 genes, about half of them can be edited by these three CRISPR nucleases (Figure 6E). PAM sequence lying in the sense (S) or the anti-sense (AS), or both sense and anti-sense (S+AS) sequences were counted respectively. The genes carrying over 50% mutations editable by either of the three CRISPR nucleases occupy half of the 20 genes, including *JAK2*, *EGFR*, *BRAF*, *IDH1*, *TERT*, *PIK3CA*, *CTNNB1*, *MUC16*, *LRP1B*, and *DNMT3A* (Figure 6F). The range of the SNV mutations that can be edited of each gene varies between 20.7% to 70.7%, and the highest predicted editing frequency is in *TERT* gene by SpCas9. What obvious is that the distribution of LbCpf1 PAM sequence is less frequent than that of SpCas9 and SaCas9. Altogether, specific targeting of cancer driver mutations by CRISPR nucleases has giant potential in treating oncogenic mutation-driven cancers, especially in the types of cancers that don’t have effective therapies. On the other hand, through bioinformatic analysis of 31555 SNV mutations, references were given to target these oncogenic mutations. At the same time, a bioinformatic pipeline was provided to analyze the distribution of PAM sequences and to estimate the target potential of other candidate genes by this high-throughput method.

## Discussion

CRISPR/Cas9 genome-editing system is a powerful technique which can specifically target genomes or their mutated sequences. In our study, CRISPR/Cas9 was used to target the *KRAS* mutant allele, but not the wild-type allele. In addition to *KRAS* mutant alleles, other cancer-driven mutations including *EGFR* mutation (L858R), genomic rearrangements (TMEM135–CCDC67 and MAN2A1–FER fusions) and *BRAF* (V600E) driver mutation were disrupted by CRISPR systems to control tumor growth^18,19,33^. Compared with *KRAS* mutations-driven cancers, which still don’t have commercially available inhibitors, there are already some EGFR inhibitors used in lung cancers with *EGFR* gene mutations, including Erlotinib (Tarceva), Afatinib (Gilotrif), Gefitinib (Iressa), Osimertinib (Tagrisso), Dacomitinib (Vizimpro) and Necitumumab (Portrazza). Clinical drugs that target cells with *BRAF* gene changes include Dabrafenib (Tafinlar) and Trametinib (Mekinist). Therefore, there is much significance in targeting *KRAS* mutant alleles, which may hold great hopes for future cancer treatment.

Compared to the traditional treatments using inhibitors of *KRAS* pathway, CRISPR-Cas system has extended the precious targeting from protein to the genomic DNA level, and this strategy can be wildly and easily spread to other oncogenic mutations. The development of traditional inhibitors, including antibodies and small molecules, is complicated and the whole process be generally designed for a single target. For example, though KRAS G12C inhibitor released by Amgen gave promising clinical outcome on its specific targets (NCT03600883), it did not show any effect on other *KRAS* mutant alleles. Besides, retargeting of different KRAS mutation at the protein level required new designment which is time- and cost-consuming. However, CRISPR system is capable to target different mutant alleles specifically and precisely at DNA level and can be easily converted to other oncogenic mutations by only changing the sgRNA sequences. On another aspect, traditional therapy could cause tumor resistance and secondary mutations, while genome-editing targets mutations at the DNA level and deplete mutations completely. Lastly, combined with NGS, individual patients can be precisely treated by CRISPR/SpCas9 targeting on their unique mutations. The editing of oncogenic mutations could also be combined with inhibitors of KRAS or other oncogenic mutations, or immunotherapy to further improve the anti-tumor efficacy.

In previous studies, CRISPR-Cas9 system was harnessed to rectify disease-associated genetic defects^34–36^ and deactivate disease-causing wild-type genes^37–39^. However, these targeting still has limited specificity without discriminating perturbation of both the wild type oncogene and mutant alleles. Our study showed that single-nucleotide mutation of a cancer driver gene in tumor cells can be selectively disrupted both *in vitro* and *in vivo* by using sgRNAs which distinguish the mutant allele from the WT one. Among the four sgRNAs targeting mutations at G12 locus, sgG12S shows the highest specificity and can discriminate a single-nucleotide polymorphism (SNP) difference in tumor cells (Figure 1B, D, E). To our best knowledge, this is the first report to demonstrate that the *KRAS* G12S mutant allele could be specifically targeted, thereby inhibiting tumor growth *in vivo*. Though Kim W. *et al*.^20^ has targeted G12V, G12D and G13D mutant alleles with lentiviral and adeno-associated viral (AAV) vectors, respectively, the mechanisms related to the tumor inhibition by targeting *KRAS* mutant alleles was not illustrated in their study. Zhao X. *et al*.^22^ has used CRISPR-Cas13a system to knockdown *KRAS* G12D allele at the transcriptional level. CRISPR-Cas13a system was reported to be tolerant to one mismatch and sensitive to two mismatches in the crRNA-target duplex, thus a second mismatch to the crRNA had to be introduced in their study, which is not so feasible to use since such a proper mutation needs to be selected out before targeting *KRAS* mutant alleles specifically. In addition, the off-target effects of the study were not assessed.

In our study, we have shown mutant allele-specific gene elimination in A549 tumors *in vivo*. Damage of the driver gene mutation *KRAS* G12S allele in A549 tumors resulted in the inhibition of cancer cell growth. Besides, on- and off-target indels as well as cell cytotoxicity associated with CRISPR/Cas9 editing were not detectably in H2228 cells which harbor wild-type *KRAS* alleles. These results are consistent with *in vivo* data that tumor growth inhibition was not observed in AdV-Cas9-sgG12S treated H2228 tumors, demonstrating the specificity of CRISPR/Cas9 for targeting a mutant allele that is in the seed sequence. The finding was in line with the previous report by Cong *et al*.^16^. In another study, CRISPR/Cas9 was used to target a mutant allele where the single nucleotide mutation generates a 5’-NGG-3’ PAM sequence that WT allele did not have, thus enabling specific targeting of mutant allele by Cas9 nuclease^18^. To extend this strategy to other cancer-driven mutations that, either locate in seed sequence or generate PAM sequences recognized by SpCas9 or other Cas9 variants, we chose top 20 mutated genes and analyzed whether their mutations could be targeted by SpCas9, SaCas9 and LbCpf1, by analyzing the seed region and PAM sequence (Figure 6E, 6F, Additional file 1: S4). We found that PAM sequence of CRISPR nucleases, especially for SpCas9 and SaCas9, are widely distributed around the mutated sites. The results indicate that this approach could be widely used to target other oncogenic mutations and could also be applied to other Cas9 families or variants. Furthermore, this approach could be utilized for multiple gene editing in cancers which are frequently characterized by mutation heterogeneity, and to test functional relevance of tumor mutations employing CRISPR/Cas9^40,41^.

In contrast to the two previous studies^20,22^, we also assessed the off-target effects *in vitro* (Figure 3A, 3B) and found there were low off-target effects during our gene targeting. Besides, we found the related mechanisms that disruption of *KRAS* G12S allele leads to blockade of AKT and ERK signaling pathways that was confirmed by WB results, thus inhibiting tumor growth. Furthermore, we assessed both the non-cutting system dCas9-KRAB and Cas9-sgG12S cleaving system and found that the transcription repression system is also capable to inhibit tumor growth both *in vitro* and *in vivo*, but at a lower efficiency. Given that dCas9-KRAB-sgG12S treatment only lead to transient transcription repression when binding rather than change the genome sequence of the mutant gene, the continuous growth inhibition of proliferating tumor cells may not be achieved completely using dCas9-KRAB-sgG12S. From this angle, the genome-editing CRISPR/Cas9 system is more practical to eliminate KRAS activation persistently. Based on our data, this mutation-sgRNA designing strategy is capable to distinguish the mutant allele from WT one at the resolution of single nucleotide differences, thus enables *KRAS* mutation-targeting at a high specificity, which is also beneficial to treat a broad spectrum of oncogenic mutations. Among thousands of mutations of the top 20 cancer driver genes we surveyed, above 50% mutations of ten genes have potential to be targeted by CRISPR system through our bioinformatic analysis. Not every oncogenic mutation can be specifically targeted due to the lack of PAM sequence, and our bioinformatic pipeline provides an easy, efficient, and high-throughput way to predict the editable sites.

## Conclusions

In conclusion, we systematically demonstrated gene-editing and mRNA-regulating systems targeted *KRAS* G12S mutant allele specifically and both *in vitro* tumor cell proliferation and *in vivo* tumor growth were inhibited. In addition, bioinformatic analysis of 31555 SNP oncogenic mutations provided a pipeline to analyze the distribution of PAM sequence for editing targets screening.

## List of abbreviations

PDACs: pancreatic ductal adenocarcinomas
CRCs: colorectal adenocarcinomas
LACs: lung adenocarcinomas
CRISPR: Clustered regularly interspaced short palindromic repeats
SpCas9: S.pyogenes CRISPR associated protein 9
EGFR: epidermal growth factor receptor
dCas9: dead Cas9
KRAB: Krüppel associated box
PAM: protospacer adjacent motif
NGS: next generation sequencing
CFA: colony formation assay
DSB: double stand break
IHC: immunohistochemical
WB: western blot
SNV: single nucleotide variation
S: sense
AS: anti-sense
SNP: single-nucleotide polymorphism
AAV: adeno-associated viral
ATCC: American Type Culture Collection
DMEM: Dulbecco’s modified Eagle’s medium
GAPDH: glyceraldehyde 3-phosphate dehydrogenase
H&E: hematoxylin and eosin

## Declarations

### Ethics approval and consent to participate

The mouse model studies were performed according to the guidelines provided by the Chinese Animal Welfare Act and approved by the Institutional Review Board on Bioethics and Biosafety of BGI.

### Consent for publication

Not applicable.

### Availability of data and materials

Data supporting this study have been deposited in the CNSA (https://db.cngb.org/cnsa/) of CNGBdb with accession code CNP0000672, and submitted to the NCBI (PRJNA576375) available online: https://www.ncbi.nlm.nih.gov/bioproject/576375.

## Funding

This work was supported by National Natural Science Foundation of China (No.81903159 and No.81502578), Science, Technology and Innovation Commission of Shenzhen Municipality (No. JCYJ20170817145218948 and JCYJ20170817150015170).

### Authors’ contributions

C-CC, YG and YM supervised the project and revised the manuscript. QG and XH designed and performed the research, and QG wrote the manuscript. WO and BK performed the research and revised the manuscript. YX designed and performed the mouse experiments. RD, YL, EW, LC, XD, YL and BZ performed the experiments. LH, DW and ZZ performed the data analyses. YH and HY reviewed the manuscript.

## Acknowledgements

We thank Yuping Ge for her support in the mouse experiments. Thank Lu Lin and Xiaoyu Wei for their support in data analysis. Thank Huanyi Chen for her help in experiments. Thank Anne Huang for revising our manuscript. Thank the support of Guangdong Provincial Key Laboratory of Genome Read and Write, and Guangdong Provincial Academician Workstation of BGI Synthetic Genomics.

## Additional file

### Additional file 1

**Figure S1.** Maps of pX330 vectors. **Figure S2.** Editing efficiency and inhibition of tumor cells of AdV-Cas9-sgG12S adenovirus. **Figure S3.** Tumor weights of xenograft mice treated with CRISPR system. **Figure S4.** PAM analysis of three CRISPR nucleases. **Table S1.** List of PCR primers used in targeted deep sequencing.

**Figure S1. Maps of pX330 vectors**, including pX330-U6-Chimeric blank vector and pX330-U6-sgRNA expressing vector.

**Figure S2. Editing efficiency and inhibition of tumor cells of AdV-Cas9-sgG12S adenovirus**. **a** Maps of adenoviral vectors, including AdV-Cas9 blank vector and sgG12S guide RNA expressing vector AdV-Cas9-sgG12S. **b** Gene editing efficiency and specificity of AdV-Cas9-sgG12S adenovirus were confirmed by sanger sequencing in A549 and H2228 cells. **c** Gene editing efficiency and specificity of AdV-Cas9-sgG12S adenovirus were confirmed by sanger sequencing in A549 and H2228 cells. **d** CCK-8 assay. Cell proliferation was accessed by using CCK-8 reagents at different timepoints after plating. The number of cells in cultures with different treatments was determined by the optical density at 490 nm.

**Figure S3. Tumor weights of xenograft mice treated with CRISPR system**. **a** Body weights of euthanized A549 tumor-bearing mice treated with PBS, AdV-Cas9 and AdV-Cas9-sgG12S on 28 days post adenoviral injection, and **b** Body weights of euthanized H2228 tumor-bearing mice on 7 days post adenoviral injection of PBS, AdV-Cas9 and AdV-Cas9-sgG12S. **c** Body weights of euthanized A549 tumor-bearing mice on 28 days post lentiviral injection of PBS, V2 and dCas9-KRAB-sgG12S, and **d** Body weights of euthanized H2228 tumor-bearing mice on 7 days post lentiviral injection of PBS, V2 and dCas9-KRAB-sgG12S.

**Figure S4. PAM analysis of three CRISPR nucleases**. **a** Top, appearance of SpCas9 PAM sequence in the sense strand of oncogenic mutations. Only when the mutation occurs in the seed sequence or PAM sequence, it can be specifically targeted. But if the mutation occurs in the N of PAM NGG sequence, it can’t be targeted specifically. This situation is considered meaningless. M, mutation, in red. Green arrow, the direction of PAM shift. Bottom, appearance of SpCas9 PAM sequence in the anti-sense strand of oncogenic mutations. **b** PAM analysis of SaCas9 in the sense and anti-sense strands of oncogenic mutations. PAM sequence of SaCas9 is NGRRN, if the mutation occurs at any N of the PAM sequence, this situation is meaningless. M, mutation, in red. Green arrow, the direction of PAM shift. **c** PAM analysis of LbCpf1 in the sense and anti-sense strands of oncogenic mutations. PAM sequence of LbCpf1 is TTTV, V is all but T. If the original V is T, then the mutation of V could lead to the specific targeting. M, mutation, in red. Green arrow, the direction of PAM shift.

**Table S1. List of PCR primers used in targeted deep sequencing.**

