## Additional file 1 for "Selective targeting of an oncogenic *KRAS* mutant allele by CRISPR/Cas9 induces efficient tumor regression": Supplemental Table 1-GM.pdf

Table 1 List of PCR primers used in targeted deep sequencing

| No. | Forward (5' to 3') | Reverse (5' to 3') |
|---|---|---|
| OnT | CTGGTGGAGTATTTGATAGTGTA | ATTCGTCCACAAAATGATTCTGA |
| OT1 | AATCC CAGCCCACTGCTTTGAG | AATCCCTCCCCAGCACCCG |
| OT2 | TGGTTATGTTTCCTTTTTGACTGC | CCCTGGAGATTCTGATACAGTGGA |
| OT3 | GCACAGAAGAACAGCAGCGAGGTAG | CCATAAAAAATCTCATCAGCCCCAA |
| OT4 | TTGTCTCTGCCCTTATGGATTGC | CCAAGAGAAAAGCATTTGTCCTGAG |
| OT5 | GCAGGATGTAGATGTGGGTAAGG | ATTAGGTGGTAAAGGTCGTGGGAA |
| OT6 | CAGGTGAAAGAAATCGTTAGGGACA | ACAAACAGTTCCTGGCACACG |
| OT7 | AGGTTATGTGGCTTATTCAAGGTCA | GTCGTAACCTTCTATGTGACTATTG |
| OT8 | CCTGAAGAGATGGGTGTATTTTGG | TAAGCCTTTCTACCTCCTGGGG |
| OT9 | CCACAAAGAAATGAGAAACAGT | TAACCCCTAAGTATTATCCCAGAA |
| OT10 | GTCACACAGTCAGTGGCAGAGAAGA | ACAGTACAGGCACAGGGTGGCAGC |
| OT11 | GTGAGGCAAGGAAGTTTGATTTT | TTCTCTATCTCCAGTCTCTGCTTT |
| OT12 | GGATAAGAGCACTTGGGCAGA | AACTGTAGGAAATAAGCAAAGGAGA |
| OT13 | CACACAAAATCCATAAAATCGGTCA | GCTACAAGAATGGTACACCCAGTCG |
| OT14 | CTGAACTAAAGAAGGAACTGCCGCC | ACTCCCGTCGTTCGGCTCGGTCCT |
