## Supplementary figures and images for "Selective targeting of an oncogenic *KRAS* mutant allele by CRISPR/Cas9 induces efficient tumor regression"

### Supplemental-1,2-GM.pdf

Figure S1

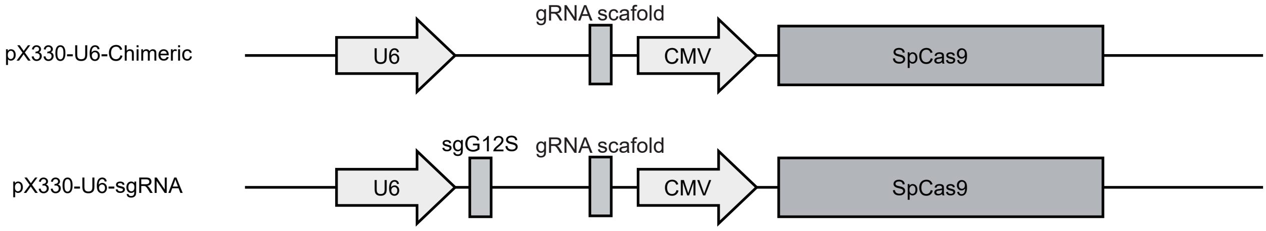


Figure S2

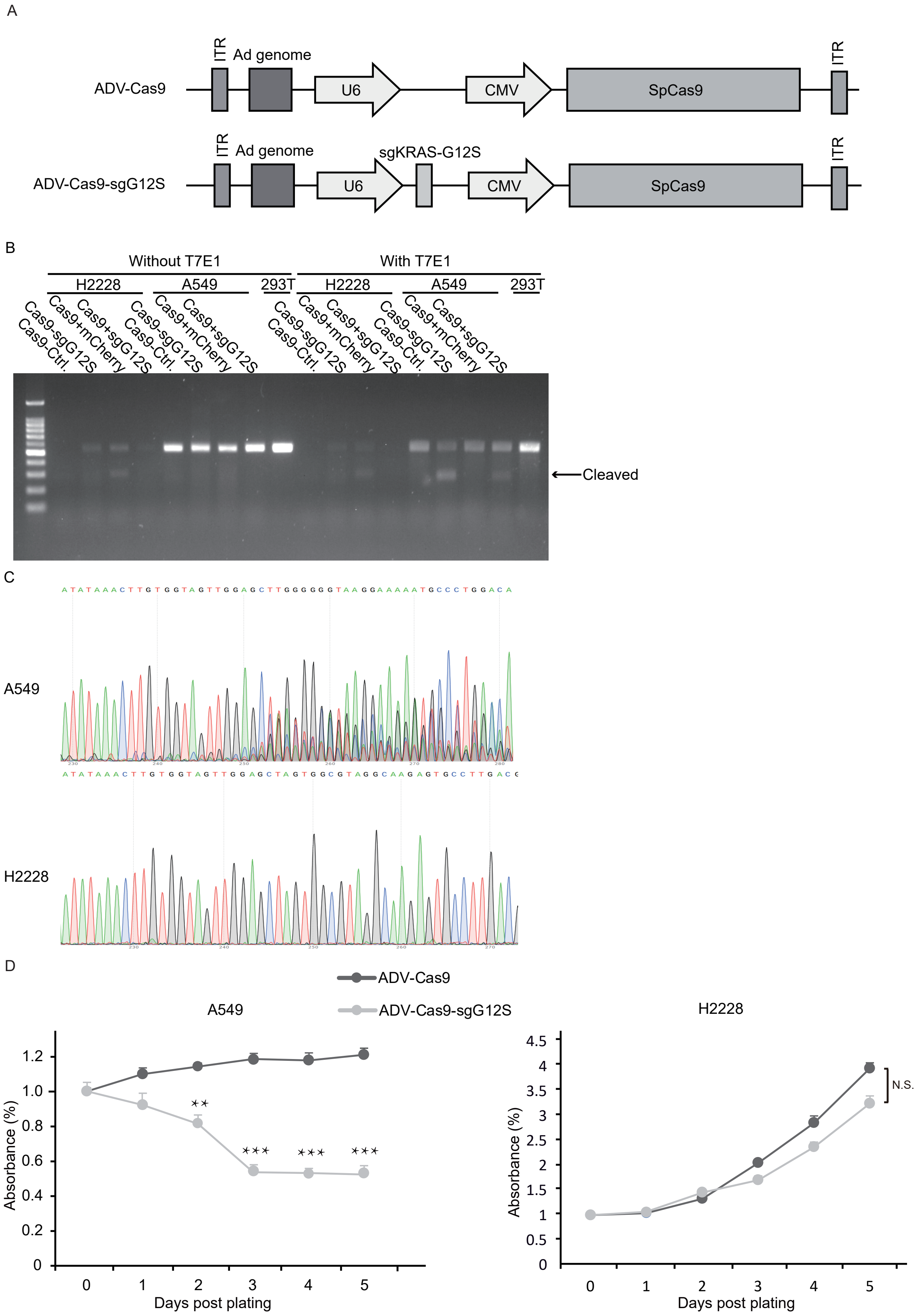

### Supplemental-3-GM.pdf

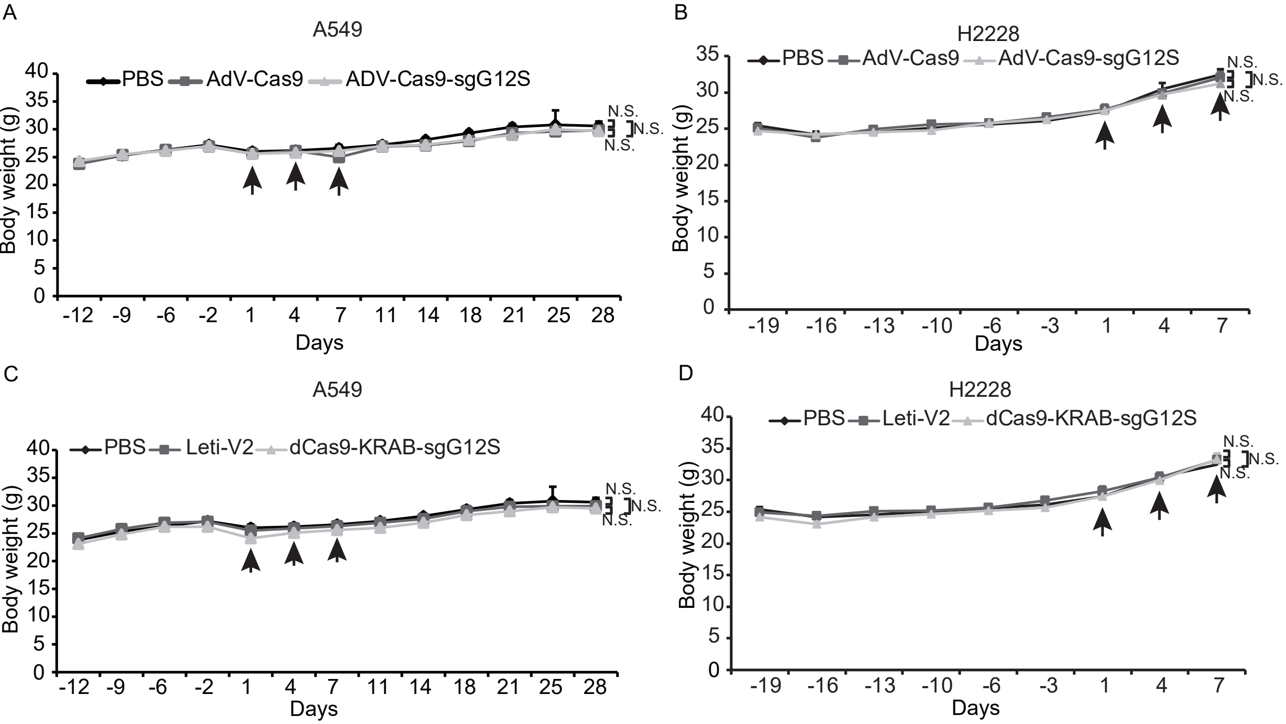

### Supplemental-4-GM.pdf

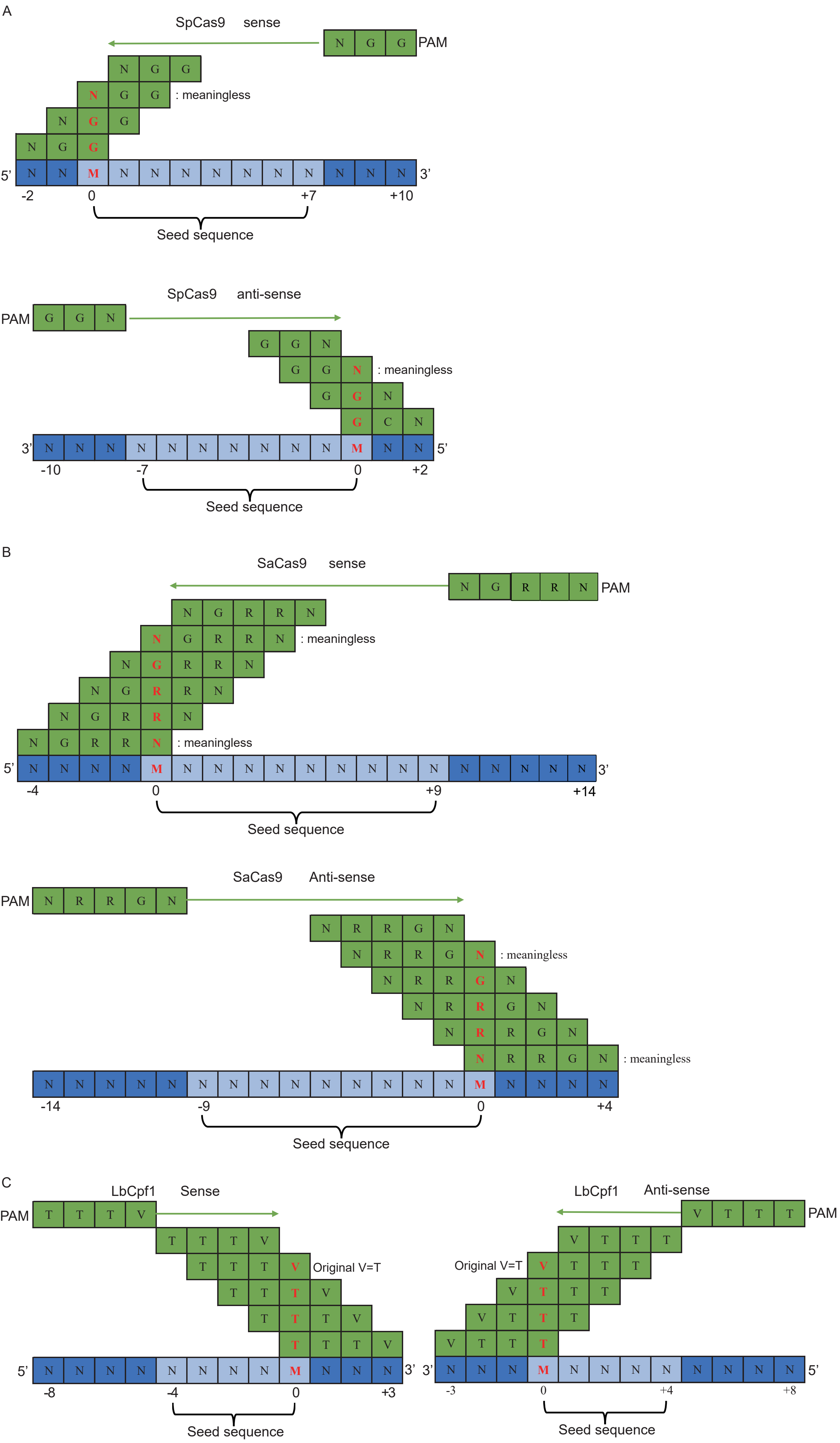
